# SASH1 interacts with TNKS2 and promotes human melanocyte stem cell maintenance

**DOI:** 10.1101/2023.09.26.559624

**Authors:** Karoline A. Lambert, Christopher M. Clements, Nabanita Mukherjee, Theresa R. Pacheco, Samantha X. Shellman, Morkos A. Henen, Beat Vögeli, Nathaniel B. Goldstein, Stanca Birlea, Jennifer Hintzsche, Aik-Choon Tan, Rui Zhao, David A. Norris, William A. Robinson, Yizhou Wang, Jillian G VanTreeck, Yiqun G. Shellman

**Affiliations:** Department of Dermatology, University of Colorado Anschutz Medical Campus, School of Medicine, Aurora, CO 80045; Department of Computer Science, University of Colorado Boulder, Boulder, CO 80309; Department of Biochemistry and Molecular Genetics, University of Colorado Anschutz Medical Campus, School of Medicine, Aurora, CO 80045; Charles C. Gates Regenerative Medicine and Stem Cell Biology Institute, University of Colorado Anschutz Medical Campus, School of Medicine, Aurora, CO 80045; Next Level Bioinformatics LLC, Dixon, IL 61021; Huntsman Cancer Institute, Departments of Oncological Sciences and Biomedical Informatics, University of Utah, Salt Lake City, UT 84112; Division of Medical Oncology, University of Colorado Anschutz Medical Campus, School of Medicine, Aurora, CO 80045; Department of Chemistry, Emory University, Atlanta, GA 30322; College of Biological Sciences, University of Minnesota, Twin Cities, St. Paul, MN 55108

## Abstract

Both aging spots (hyperpigmentation) and hair graying (lack of pigmentation) are associated with aging, two seemingly opposite pigmentation phenotypes. It is not clear how they are mechanistically connected. This study investigated the underlying mechanism in a family with an inherited pigmentation disorder. Clinical examinations identified accelerated hair graying and skin dyspigmentation (intermixed hyper and hypopigmentation) in the family members carrying the SASH1^S519N^ variant. Cell assays indicated that SASH1 promoted stem-like characteristics in human melanocytes, and SASH1^S519N^ was defective in this function. Multiple assays showed that SASH1 binds to tankyrase 2 (TNKS2), which is required for SASH1’s promotion of stem-like function. Further, the SASH1^S519N^ variant is in a *bona fide* Tankyrase-binding motif, and SASH1^S519N^ alters the binding kinetics and affinity. Results here indicate SASH1 as a novel protein regulating the appropriate balance between melanocyte stem cells (McSC) and mature melanocytes (MCs), with S519N variant causing defects. We propose that dysfunction of McSC maintenance connects multiple aging-associated pigmentation phenotypes in the general population.

## INTRODUCTION

Melanocytes (MCs) are specialized pigment-producing cells responsible for human skin and hair pigmentation. Dysregulation of MCs leads to diseases impacting general health and quality of life, most notably melanoma, but also common pigmentation abnormalities such as lentigines ^1^. However, little is known about the molecular mechanisms for controlling McSC and MC, or mechanisms behind many pigmentation disorders. Lentigines are small, hyperpigmented skin spots, histologically containing an increased number of MCs with elevated amounts of melanin ^2^. Solar lentigines are common, acquired pigmented lesions, also known as sunspots, age spots, or liver spots ^3,4^. Histologically, aging spots exhibit increased numbers of MCs in the epidermis ^5^, and are common on sun-exposed areas of the skin, such as the face and hands ^6-8^. The molecular etiology of aging spots is not well defined. Hair graying, that is a lack of pigmentation in the hair shaft, is the hallmark of aging in mammals ^9^. It is intriguing that both increasing-pigmentation and lacking-pigmentation are associated with aging, and it is unclear whether they share the same pathological mechanisms. We present a family with a monogenic pigmentation disorder carrying the SASH1^S519N^ variant, which connects these two seemingly opposite aging-associated phenotypes. Monogenic inherited disorders provide unique human transgenic-like models to uncover the molecular mechanisms involved in complicated biological processes.

SASH1 is a scaffold protein with context-dependent biological functions in cell adhesion, tumor metastasis, lung development, and pigmentation ^10^. We previously identified a heterozygous variant, SASH1^S519N^, as causative for inherited lentigines in a Hispanic family in which lentigines appear in children (< 10 years old) ^11^. All individuals with the variant are affected and display similar skin pigmentation phenotypes without any other obvious abnormalities ^11^. Additional SASH1 variants have been associated with similar inherited, autosomal dominant pigmentation disorders ^11-23^. They are diagnosed as either multiple lentigines or Dyschromatosis Universalis Hereditaria, which is a rare genodermatosis characterized by the co-existence of hyper- and hypo-pigmented macules throughout the whole body. Here, we investigated the underlying mechanism for the S519N family, and the results identify novel proteins regulating McSC maintenance and provide insights into the development of age spots and gray hair in the general population.

## RESULTS

### SASH1^Nonvariant^, but not SASH1^S519N^, promotes stem-like characteristics in human MC cultures

Lentigines of SASH1^S519N^ (S519N) individuals show an increased population of skin MCs ^11^, which could develop via several mechanisms, including increased proliferation or survival of McSCs and/or MCs. Without the availability of stable human McSC cell cultures, we employed a sphere-forming assay with human primary MCs; the sphere assay selects for stem-like characteristics ^24^. Compared to the empty vector (EV) control, expressing SASH1^Nonvariant^ (N.V.) — but not S519N — significantly increased the size and number of formed spheres (**Fig. 1A and C**). Results were validated with two different expression vectors (HA-tagged or GFP-tagged), and **Fig. 1A** summarizes five independent experiments. We further validated these findings with the well-established Aldefluor assay (**Fig. 1B**), which measures the stem/progenitor marker ALDH+ ^25^. **Fig. 1D** shows the similar expression levels of SASH1 between N.V. and S519N in the MCs. Conversely, knocking down SASH1 in MC cells with siRNA decreased the number of formed spheres (**Fig. S1**). These results indicated that N.V., but not S519N, promotes stemness of human McSC.

**Figure 1.**
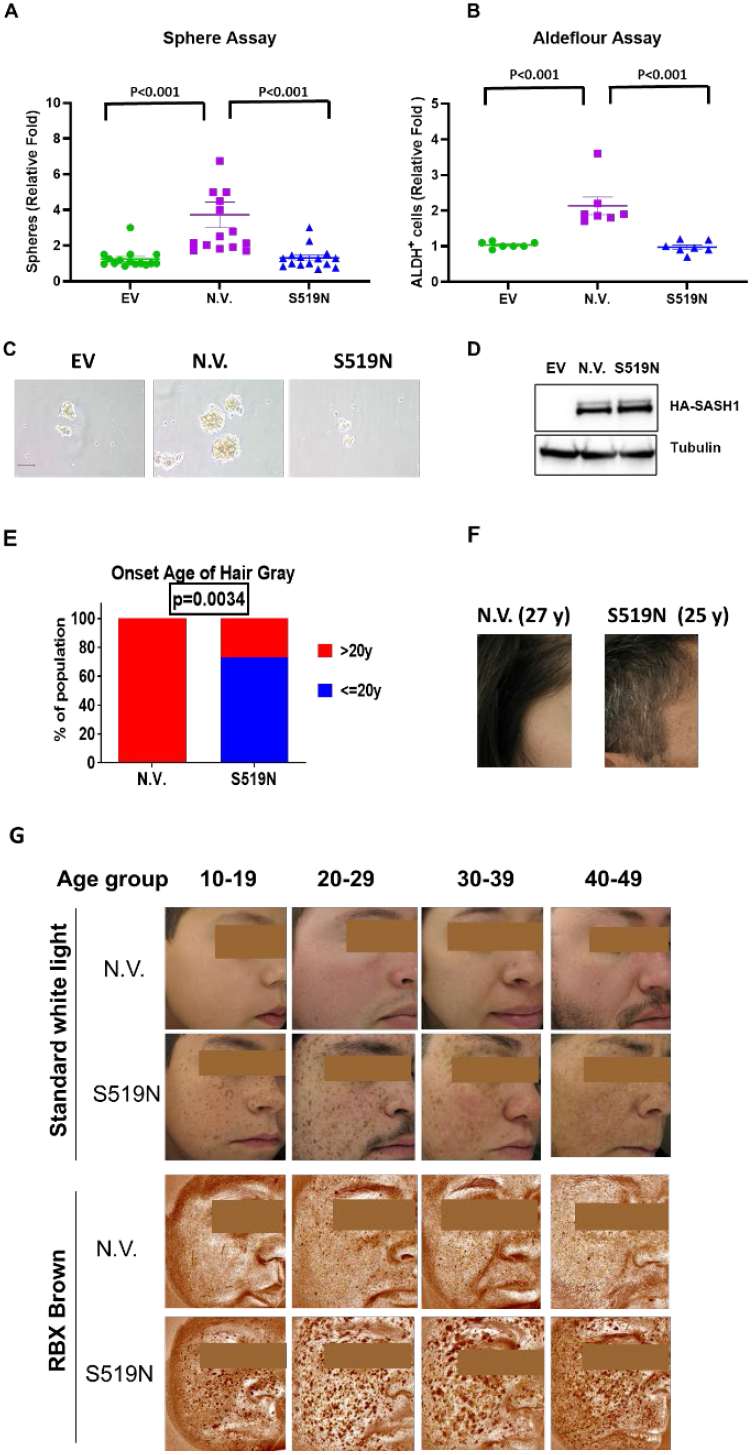
Impact of SASH1 (N.V. and S519N) on McSC functions in cells, in human hair and in skin pigmentation. Human primary epidermal melanocyte cells (HEMn-MP) were transfected with control vector (EV),N.V. SASH1 or S519N. Cells were then subjected to the sphere forming assay (**A** and **C**), Aldefluor assay (**B**), or immunoblot (**D**). **E**. Early onset of hair-graying in the S519N individuals. % members from N.V. (0/8) or S519N family members (8/11) with onset of hair-graying at *≤* or > 20 years old. **F**. Examples of hair in an individual carrying N.V. or S519N of SASH1. **G**. The standard white light images and melanin pigmentation of RBX®-Brown cross-polarized images of indicated individuals by age range. Images captured with a VISIA-Complexion Analysis multi-modality facial imaging system.

### SASH1^S519N^ accelerates the exhaustion of McSC pools in humans, as seen by early onset of gray hair and skin dyspigmentation

The decrease of McSC stemness leads to McSC exhaustion, with gray hair as the hallmark ^9^. Previous studies estimate that 41.6 years is the average age for the onset of gray hair in the general population ^26^. Strikingly, SASH1^S519N^ individuals show early hair graying as young as age 11, and 73% (8 out 11) reported the onset of gray hair by the age of 20, significantly different from SASH1^N.V.^ family members (p = 0.0034) (**Fig. 1E)**. Four individuals reported gray hair before age 15, and two had completely white hair by the age of 30. In contrast, no N.V. SASH1 family members (n = 8) had gray hair younger than 20 years old. **Fig. 1F** is an example of an SASH1^S519N^ individual with salt-pepper hair at 25 years old, compared to a SASH1^N.V.^ family member at 27 years old. It turns out that many affected individuals began dyeing their hair at a young age to conceal the early onset of gray hair.

One would predict that exhaustion of the McSC pool leads to eventual skin hypopigmentation when mature MCs in skin are replaced during normal skin maintenance. Here, we further examined the facial pigmentation phenotypes using the VISIA multi-modality facial imaging system, including non-variant family members of similar ages as controls (**Fig. 1G**). The skin of non-variant individuals displayed even color tone, in both standard light photography and RBX®-Brown cross-polarized images, which shows melanin below the skin surface that is not visible to the naked eye. In contrast, there was an obvious increase in both hyperpigmentation and hypopigmentation in the facial skin of individuals with the S519N variant, and hyperpigmentation also diminished with increased age (**Fig.1G**). The SASH1^S519N^ individuals exhibited notably higher levels of hyper- and hypopigmentation compared to their age-matched family members, and dyspigmentation appeared at a young age. Co-mingling of hyper-pigmented and hypo-pigmented patches on the skin is defined as dyspigmentation, which is a photoaging phenotype increasing with age ^27^ in the general population. The SASH1^S519N^ phenotypes have remarkable similarities to the trajectory of skin dyspigmentation and the loss of hair pigment, simply beginning at a younger age (see the proposed model in **Fig. 2A**). Thus, the pigmentation phenotypes in SASH1^S519N^ individuals are also consistent with the concept that the SASH1^S519N^ disorder may represent an example of accelerated aging of human McSC.

**Figure 2.**
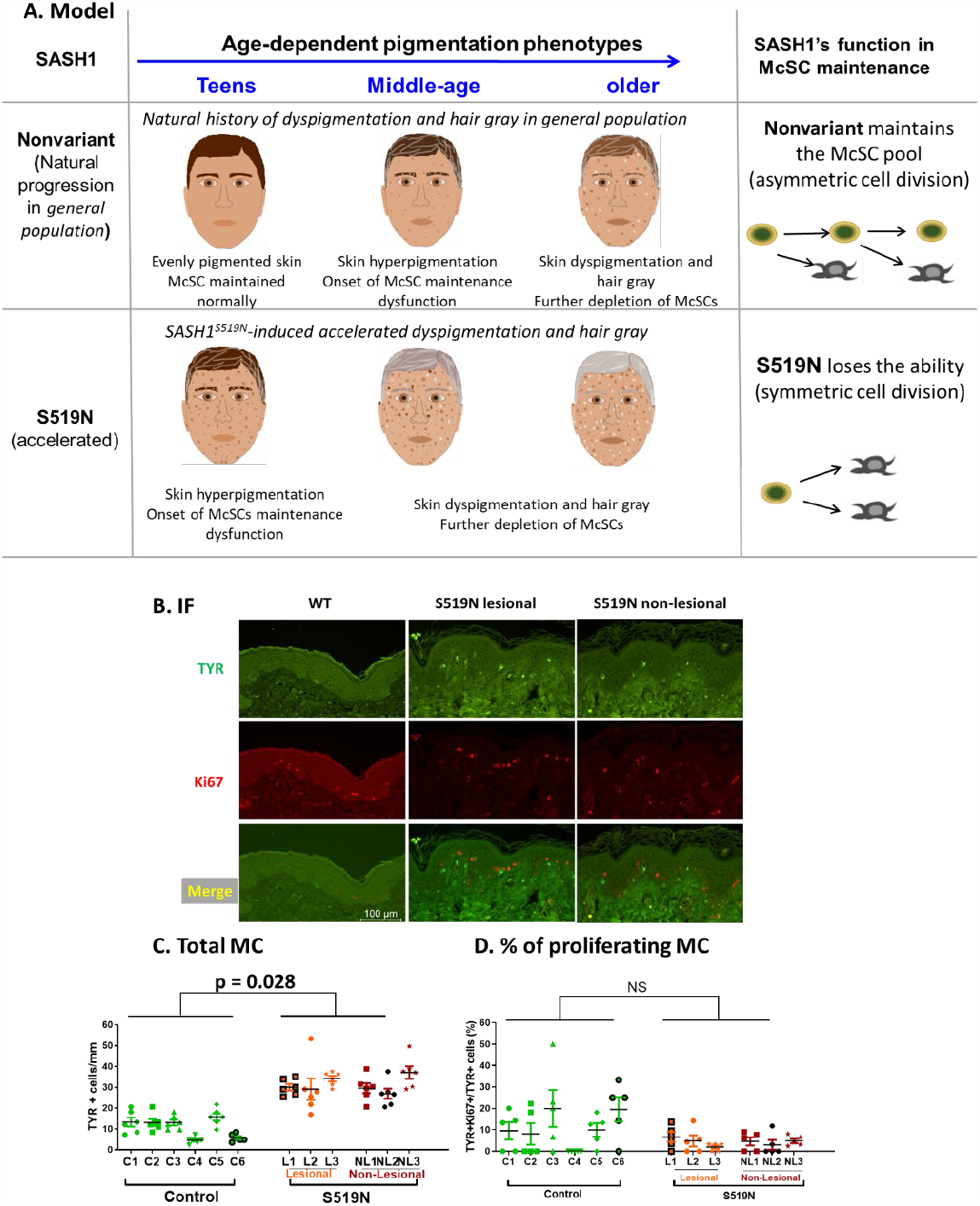
SASH1^S519N^ increased non-proliferating MC numbers in the skin. **A** shows the proposed model of accelerated pigmentation phenotypes and SASH1’s function in McSC maintenance. In the general public, middle-age brings the onset of lentigines and gray hair, but SASH1^S519N^ individuals develop by their teens. **B**. Immunofluorescent (IF) staining of skin biopsies with the MC marker (TYR in green) and a proliferation marker (Ki67 in red). Skin biopsies were collected from unaffected family members or non-related control individuals (control) and from individuals carrying S519N. Lesional (hyper-pigmented) and adjacent non-lesional areas were stained. Scale bar = 100 um. **C&D** show quantification, with the mean for each group at the bottom. The quantification of the TYR or Ki67 staining was performed by counting the number of positively stained cells per millimeter of tissue across the entire length of the biopsy. **C** shows the number of MCs in skin (TYR+ cells), and **D** shows % of proliferating MCs (TYR+KI67+/TYR+), with 6 in the control group and 3 in the S519N group.

### The skin of SASH1^S519N^ individuals displayed an increased number of non-proliferating MCs

To examine the effects of S519N on the proliferation of MCs in human skin, we performed co-immunofluorescent staining with the markers for differentiated MCs (Tyrosinase, TYR) and proliferation (Ki67) of skin biopsies from N.V. and S519N individuals of ages between 20–50 years (**Fig. 2B**). The number of differentiated MCs (TYR+ cells) were 2-3 fold higher in S519N, in both lesional (hyper-pigmented) and non-lesional (adjacent non hyper-pigmented) sections (**Fig. 2C**), consistent with our previous finding ^11^. However, the percentage of proliferating MCs (TYR+Ki67+ double positive cells) was not increased (**Fig. 2D**). These data suggest that this larger MC population is not due to hyper-proliferation, a proposed mechanism for other well-known inherited lentigines ^1^. These results are also consistent with our cell culture studies and support the concept that SASH1^S519N^ disrupts the appropriate McSC maintenance and shifts the balance toward mature MCs.

### SASH1 binds to TNKS2, and TNKS2 is required for SASH1’s function

To investigate the functions of SASH1 at the molecular level, we used yeast-two-hybrid (Y2H) screens to identify the binding partners for SASH1. Tankyrase 2 (TNKS2) is a top candidate (**Fig. 3A**). Tankyrases belong to the multifunctional poly(ADP-ribose) polymerase family that modify proteins through ADP-ribosylation, and tankyrases also act as protein scaffolds ^28,29^. Tankyrases play crucial roles in multiple cellular functions and disease pathologies, through altering the activity, stability, and/or subcellular location of their binding partners ^28,29^. All their binding partners have at least one Tankyrase-binding-motif (TBM), which binds to one of the Ankyrin Repeating Cluster (ARC) domains in the tankyrases ^30^. The canonical TBM’s sequence is RxxxxG-[no P]-x ^30^. Interestingly, S519 is located in a predicted TBM (**R**^512^SSLS**G**QS^519^) (**Fig. 3B**), and we found nine potential TBMs in SASH1 (**Fig. 3C**), using our bioinformatics tool (manuscript submitted). These imply that the interaction between SASH1 and TNKS2 is part of the etiology of this disorder.

**Figure 3.**
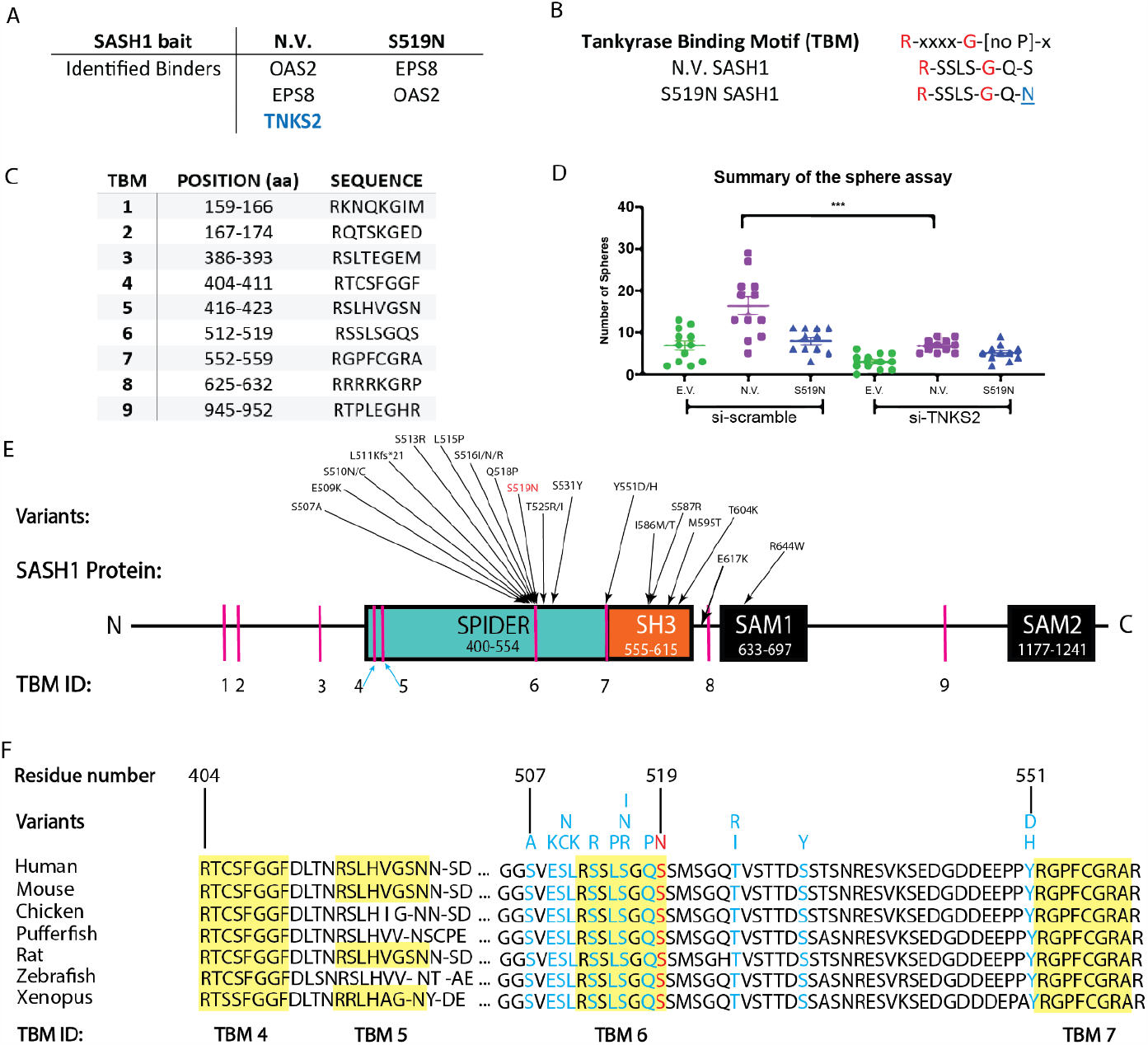
TNKS2 binds to SASH1 and is required for SASH1’s function. **A**. The top binders to SASH1 were identified from Y2H screens using full length human SASH1 (N.V. or S519N) as a bait and a cDNA library generated from human MCs as the prey. **B**. S519N locates in a predicted canonical TBM. **C**. Nine predicted TBMs in SASH1. **D**. Human primary MCs were transfected with expression vectors of SASH1 (control vector, EV; Non-variant, N.V. or S519N SASH1), with siRNAs si-Scramble control or against TNKS2. Cells were then subjected to sphere forming assays. *** indicates p < 0.0001. 60-70% knockdown was achieved (Fig.S2). **E**. Locations of the TBMs and the variants associated with pigmentation disorders in SASH1 show most of these variants locate within or close to a TBM. **F**. The SPIDER region is conserved within vertebrates, contains four TBMs (highlighted sequences) and contains most hyper-pigmentation variants, including S519N in TBM 6.

To determine if TNKS2 affects SASH1’s function in stem-like features of the MC lineage, we performed similar sphere assays as in **Fig1.A** with human primary MCs and with siRNAs, either the scramble control (si-Scramble) or those against TNKS2 (si-TNKS2) (**Fig.3D**). Consistent with **Fig1.A**, N.V., but not S519N, promoted sphere formation. Strikingly, knocking down TNKS2 severely diminished the enhanced sphere-forming capacity of N.V. SASH1. These results demonstrate that TNKS2 is required for SASH1’s function in promoting stem-like characteristics in the MC lineage.

S519 is located in a highly conserved region of SASH1 (amino acid 404-560), which contains four putative TBMs (**Fig. 3E**). Importantly, this region is central for SASH1’s roles in pigmentation, which contains ∼70% of the SASH1 variants (17 out of 24 so far) associated with pigmentation disorders ^11-23^. Further, at least 16 variants are located within, or close to, the TBM6 (aa 512-519) and TBM7 (aa 552-559) (**Fig. 3E-F**). Because this region (aa 400-554) is highly conserved and structurally disordered, we refer to it as SPIDER (<u>S</u>Ly <u>p</u>roteins associated disordered region) ^31^. These results implicate the importance of the interaction between SASH1’s SPIDER and TNKS2 in human pigmentation.

### S519N locates in a bona fide TBM and alters how TNKS2 binds to SASH1

To determine the effects of the S519N variant, we performed co-IP experiments (**Fig. 4A)**. Both N.V. and S519N SASH1 bind to TNKS2, indicating that S519N did not abolish the binding between SASH1 and TNKS2. To better characterize their binding, we then utilized NMR spectroscopy to determine the regions of direct interaction in SPIDER and assess their binding kinetics. Due to the disordered nature of SPIDER, NMR is the best-suited technique for its atomic level characterization. Two-dimensional fingerprint ^15^N-^1^H HSQC and 1D proton NMR spectroscopy showed SPIDER of N.V. SASH1 is disordered (**Fig. 4B**), while ARC4 is well folded (**Fig. S3**) as reported in ^32^. Overlay of N.V. and S519N SPIDER ^1^H-^15^N HSQC spectra indicated that the variant does not result in significant structural changes in free forms ^31^. Next, we monitored changes in the peaks of SPIDER as a function of titrating the ARC4 domain of TNKS2. The changes can be chemical shift perturbations (CSPs), indicating a change in the chemical environment, or peak broadening (and disappearance), which could be due to an increase in overall tumbling of the formed complex or micro- to millisecond binding dynamics. Indeed, titrations of unlabeled ARC4 into 100 µM of ^15^N-SPIDER ranging from 0 to 1.2 mM show peak broadening and significant CSPs indicative of binding (**Fig. 4B**). At the 1:4 titration point, we observe significant CSPs within the predicted TBMs 4, 5, 6, and 7 (**Fig. 4C**). Thus, SASH1 contains *bona fide* TBMs, and S519N is in one of them.

**Figure 4.**
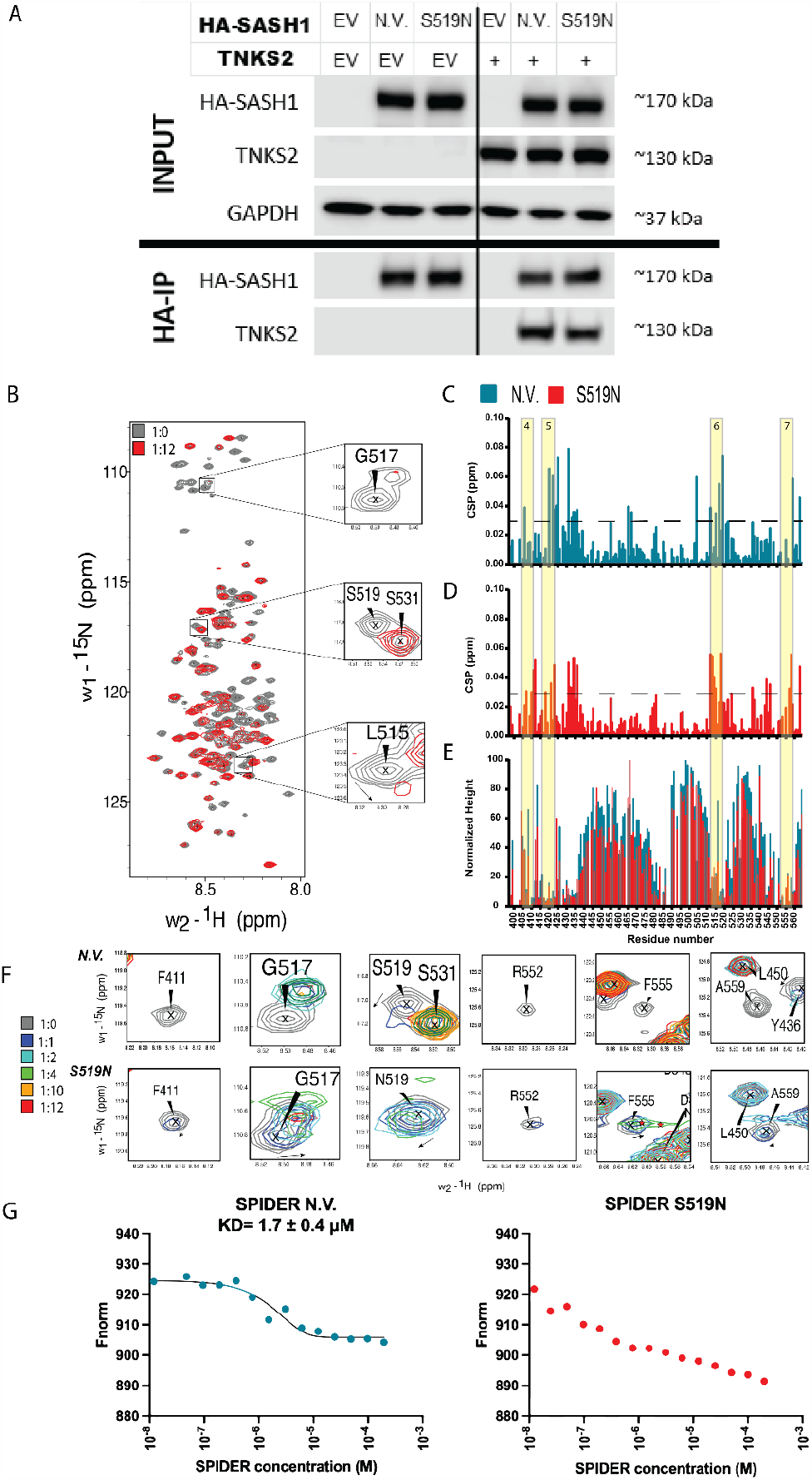
S519N variant alters the dynamic, kinetic and affinity of the binding between SASH1 to TNKS2. **A**. HA-tagged SASH1 (Empty vector (EV), N.V. or S519N) were co-transfected with FLAG-tagged TNKS2 (EV or TNKS2) in 293FT cells. Total lysates (INPUT panels) showed the protein expression. Co-IP with HA-tagged magnetic beads showed robust pull-down of TNKS2 by SASH1 (HA-IP panels). **B**. ^15^N-^1^H HSQC spectrum of human non-variant SASH1-SPIDER (<u>396-564</u>) (gray) is shown overlaid with the 1:12 titration mark with TNKS2-ARC4 (red). Peaks upfield of 8.0 ppm are not shown. Three backbone chemical shifts from TBM 6 are expanded to show chemical shift perturbations (CSPs) (L515) or peak broadening (G517, S519). **C**. and **D**. CSPs for N.V. and S519N SASH1-SPIDER, respectively, induced by 1:4 SASH1:TNKS2-ARC4. The yellow bars indicate predicted TBMs 4, 5, 6, and 7. The dotted lines represent the averages plus one standard deviation at 0.031 and 0.029 ppm. Higher CSPs indicate a greater change in the chemical environment. **E**. Overlay of normalized peak intensity for both N.V. and S519N SPIDER at 1:12 compared to free. Lower normalized values indicate a greater reduction in intensity at the 1:12 point, thus representing a greater change in chemical environment. **F**. Select CSPs show that N.V. peaks disappear earlier than in S519N during titration. F411, G517, R552, F555, and A559 all broaden and disappear at the 1:1 titration point, whereas they all broaden at a higher titration point in S519N. Despite these differences, both S519 and N519 experience CSP and peak broadening indicating that binding is maintained at TBM 6 for both constructs. **G**. ARC4 binding affinity to SPIDER N.V. and SPIDER S519N measured by MST. Sixteen serial dilutions of SPIDER were mixed with fluorescently labeled ARC4. The *K*_D_ value for the SPIDER N.V. is 1.7 ± 0.4 μM. *K*_D_ value for SPIDER S519N could not be estimated due to weak binding.

To determine the atomic level changes of SPIDER in binding induced by the S519N variant, we repeated these titrations with S519N. Again, we observed significant CSPs at the 1:4 titration points in all four predicted TBMs (**Fig. 4D**). At the 1:12 titration, both N.V. and S519N showed a significant reduction of peak intensities in the four TBMs, compared to free SPIDER (**Fig. 4E**). Additionally, discrepancies in peak chemical shifts and intensities of N.V. and S519N reveal atomic level differences between the two constructs. A representative sample of overlaid peaks demonstrates that the N.V. peaks disappear at lower titration points in comparison to S519N, which suggests changes in binding kinetics and possibly tighter binding of N.V. (**Fig. 4F**). The highest titration points did not reach saturation, thus preventing us from fitting exact *K*_*D*_ values. Nevertheless, the quality of the titration profiles was sufficient to be used for comparative purposes. We conclude that the binding affinity for the N.V. is attenuated by approximately one order of magnitude for the variant (**Fig. S4**). The distinct differences in the positions and intensities of the peaks observed in the two constructs demonstrate that while the S519N substitution in SPIDER does not eliminate binding with ARC4, it does affect how they bind.

Due to the peak disappearance in the NMR titration, we performed microscale thermophoresis analysis (MST) to quantify the binding affinity between SPIDERs (N.V. and S519N) and ARC4 (**Fig. 4G**). From MST, we obtained a *K*_D_ value of 1.7 ± 0.4 μM for SPIDER N.V. binding to ARC4. On the other hand, we are not able to estimate an accurate *K*_D_ value for the interaction between SPIDER S519N and ARC4 due to the weaker binding and absence of saturation. These findings collectively demonstrate that the S519N substitution alters the binding kinetics of SPIDER with ARC4 and results in weaker affinity.

## DISCUSSION

The etiology of many pigmentation disorders is unclear. This study uncovered the pathological mechanism for SASH1^S519N^ hyper-pigmentation and investigated the functions of SASH1 in the MC lineage. Multiple lines of evidence, including both cell culture assays and clinical observations, reveal SASH1 and TNKS2 as novel proteins promoting human McSC maintenance. The SASH1^S519N^ family connects the early onset of both lentigines and hair graying to the dysfunction of McSC maintenance, providing a new model for studying human McSC/MC biology.

Dysfunction in McSC maintenance of the SASH1^S519N^ individuals connects multiple aging phenotypes. The SASH1^S519N^ disorder represents an example of accelerated aging of human McSCs, connecting multiple aging-associated pigmentation phenotypes: both the loss and gain of pigmentation can be associated with dysfunction in McSC maintenance. Results here have broader implications on molecular mechanisms of aging in the general population, such as aging spots, loss of hair pigmentation, and dyspigmentation. The pigmentation phenotypes in other aging-related diseases support this concept. For example, patients with progeroid syndromes, such as Hutchinson-Gilford Progeria and Werner syndrome, also display premature hair graying and co-existing of hyper and hypopigmentation in skin ^33-36^. Taken together, it is conceivable that physiological aging of the McSC pool contributes to the cause of solar lentigines in the general population (**Fig. 2A**).

We showed that SASH1 promotes stem-like feature in the MC lineage, and this function requires TNKS2. To the best of our knowledge, this study is the first to report the roles of SASH1 in stem cell maintenance. Few studies have investigated the functions of TNKS2, with most tankyrase research focusing on TNKS2’s homolog tankyrase 1 (TNKS). TNKS and TNKS2 have overlapping functions and loss of both in mice is embryonic lethal ^37^. Still, the role of TNKS in stem cells is not well understood, with various studies suggesting tissue- or stage-specific roles. For example, TNKS promotes adult intestinal stem cell proliferation in Drosophila ^38^ and regeneration of adult fin following injury in zebrafish ^39^. However, tankyrase inhibition, in combination with MEK/ERK and GSK3 inhibition, promotes the reversion of human induced pluripotent stem cells to a stable human preimplantation inner-cell-mass like naïve state ^40^.

It is unclear how SASH1 and TNKS2 promote McSC maintenance and keep a proper balance between self-renewal and differentiation. Data suggests that SASH1 and TNKS2 play a role in regulating asymmetric cell division (ACD) of McSC. The key principle of ACD is to generate distinct daughter cell fates through mechanisms linked to mitosis ^41,42^. ACD of stem cells is a highly conserved and tightly regulated process to maintain renewal capacity and tissue functions during development and tissue homeostasis ^41,42^. It produces two daughter cells with distinct fates, one stem cell and one differentiated (mature) cell, whereas symmetrical cell division (SCD) produces two daughter cells of the same fate. Shifting stem cells from ACD to SCD alters the balance between self-renewing and differentiation. It leads to diseases such as cancer when dysfunction results in two stem cells ^42^, or congenital hydrocephalus when dysfunction results in two mature cells ^43^.

SASH1 seems to play a role in mitosis and mitotic spindle function, based on our multiple pathway enrichment analyses (**Fig. S5)**. First, the mitotic cell cycle was a top enriched pathway for proteins interacting with SASH1 (**Fig. S5A**). Here, we combined the gene lists of three Y2H screens for SASH1-binding proteins (ours with two additional studies ^44,45^), and performed pathway enrichment analyses using metascape.org as in ^46^. Second, our RNAseq analysis suggests that “G2/M CHECKPOINT” and “MITOTIC SPINDLE” were the top pathways affected by knocking down SASH1 in human MCs cultured in sphere forming conditions (**Fig. S5B**). Similar effects of SASH1 knockdown on the gene expression were also seen in human vascular cells ^47^. Thus, our pathway enrichment analyses for both the SASH1’s binders at the protein level and SASH1-affected gene expression at the mRNA level, support that SASH1 has a role in mitosis and mitotic spindle. Notably, TNKS also has functions in mitosis, by promoting proper spindle assembly, structure and functions ^48-52^.

Our data (**Fig. 1**) indicate a fate shift of McSCs towards MCs in SASH1^S519N^ individuals, which resembles a change in cell division of stem cells from asymmetric to symmetric, resulting in both an increase of mature MCs and accelerated exhaustion of the McSC pool (**Fig. 2A**). We thus propose that SASH1 and TNKS2 play a role in regulating asymmetric cell division (ACD) of McSCs, and the S519N variant shifts ACD toward symmetrical cell division (SCD) (**Fig. 2A**). Consequently, without the generation of self-renewing McSCs, the pool disappears over time. Due to a lack of models, most research on the mechanism(s) driving the balance of ACD and SCD has been conducted on lower organisms. The SASH1^S519N^ pigmentation disorder may be one of the few reported cases of ACD dysfunction in human cells.

Plausibly, the binding of SASH1 and TNKS2 is involved in maintaining ACD of McSC through the mitotic function, and the S519N variant alters the binding kinetics and affinity (**Fig. 4)**, which may shift the mitotic outcome. Within SASH1, we found that multiple pigmentation-related variants cluster in or near TBMs. We showed that SASH1 binds to TNKS2 in a multivalent manner through an intrinsic-disordered region (IDR) in SASH1 (**Fig. 4**), and multivalent binding of IDRs are known to drive the assembly of multi-protein complexes ^53^. Both TNKS2 and SASH1 are scaffold proteins and can complex with multiple proteins. Further, the conformational changes in scaffolds are central to the regulation of signaling pathways and molecular processes, and mutations affecting these dynamic changes have been implicated in human diseases ^54,55^. Thus, we propose that during mitosis, S519N causes conformational changes of the protein complex containing SASH1 and TNKS2, which shifts the cell division from ACD to SCD.

In conclusion, the SASH1^S519N^ family mechanistically connects early onset dyspigmentation with early onset hair graying and demonstrates SASH1 as a novel protein regulating the appropriate balance between McSCs and mature MCs. Mechanistically, we propose the altered binding of TNKS2 to the S519N variant shift McSC cell divisions from generating one self-renewing McSC and one mature MC to generating two mature MCs. This mechanism of dysfunction in McSC maintenance may be generally relevant for pigmentation disorders. Additionally, dermatologists should include questions on the loss of hair pigment when assessing skin hyperpigmentation disorders, as many young patients may darken their hair with dye.

## ACKNOWLEDGEMENTS

This work was supported in part by NIH grants R01AR074420 and R03AR064555 to YGS, a pilot grant to YGS and TRP from University of Colorado Skin Disease Research Center grant (P30AR057212, DAN), R01 GM130694 to BV, 1R21 AI171827 to MAH, R35GM145289 to RZ, University of Colorado Cancer Center Support Grants P30 CA046934 to Flow Cytometry Core and P30 CA046934 to NMR Spectroscopy Core, and NIH Biomedical Research Support Shared Grant S10 OD025020-01. The authors thank Professor David Jones, the director of the NMR facility at the University of Colorado, Anschutz Medical Campus, for his continuous help. We thank many excellent works in the field and apologize for not being able to cite all of them.

## Competing interests

The authors declare that they have no competing interests.

## METHODS

### Transfections, sphere-forming, Aldefluor assay, immunoblots, and co-immunoprecipitations (co-IPs)

For sphere forming assays with SASH1 over-expression, 3.75 µg plasmids were introduced into 1.5 × 10^6^ HEMn-MP primary MC cells with a Nucleofector 4D system, using the P2 solution and program DS-137 as described ^56^. 5 hrs later, cultured cells were detached and either lysed for immunoblot or re-seeded for the sphere-forming ^24^ or Aldefluor assays ^25^. For co-IPs, 2.5 × 10^6^ 293FT cells were seeded onto 10 cm dishes and grown overnight. 3 μg total plasmids (1 μg pMEV-2HA-vectors + 2 μg pFLAG-vectors) were introduced with Effectene (1:10 μg DNA to reagent) and incubated for 24 hrs. Lysates were collected with 500 μl lysis buffer, incubated for 15 min, centrifuged at 13,200 rpm for 12 min, and incubated with pre-washed anti-HA-tag magnetic beads for 2 hrs. The beads were washed four times, and final lysates were collected with 50 μl elution buffer heated to 70° C for 15 min. The co-immunoprecipitation lysates were PAGE separated with a 4-12% gel and wet transferred overnight at 30 V. Details on cell culture medium, sphere forming assays, vectors, and antibodies were previously described ^24,25^ or detailed in the **Tables.S1-4**.

### Human subjects, facial pigmentation image capture, and Immunofluorescence staining

All studies were approved by the Colorado Multiple Institutional Review Board. All photographs, biopsies, and inquiries were obtained after informed consent. Images of the study subject were captured using the VISIA-Complexion Analysis (VISIA-CA) multi-modality facial imaging system according to the manufacturer’s instruction (Canfield Scientific, Inc) ^57^. VISIA-CA captures standard white light images and pigmentation images (RBX®-Brown cross-polarized images), based on their RBX® technology.

Immunofluorescence staining of skin biopsies were from non-photo-exposed skin of 6 unaffected or non-related controls and 3 individuals carrying SASH1^S519N^. Lesional (hyper-pigmented) and adjacent non-lesional areas were handled and stained as in ^11^, with antibodies in **Table.S1**.

### Yeast two-hybrid (Y2H) screens

Screens were performed with the ULTImate Y2HTM platform (Hybrigenics, Paris, France) on a human MC cDNA library as prey, with human full length SASH1 (aa1–1247) of N.V. or S519N as baits, as in ^45^, according to published protocols ^58^.

### Expression and purification of recombinant proteins in *E. coli*

Methods were used similarly as in ^59^. Vectors, buffers, and columns are listed in detail in **Table.S2-5**. ARC4-TNKS2, SPIDER-N.V., and SPIDER-S519N were expressed in BL21 (LEMO21-DE3) competent *Escherichia coli*. ARC4 and flanking residues (488-649) cDNA were codon optimized and purchased from GeneScript, cloned into a pGEX-6P-1 vector, and expressed as a fusion protein with an N-terminal GST-tag coupled via a PreScission protease cleavage site and a C-terminal 6x-Histidine tag. The SPIDER of SASH1 and flanking residues (396-564) constructs (both N.V. and S519N) were codon optimized and cloned into a pET-28a (+) vector and expressed as fusion proteins with N-terminal 6x-Histidine tag coupled via a Thrombin cleavage site. Transformed colonies were grown overnight on LB + antibiotic plates (**Table S1**) and transferred to LB + antibiotic for non-isotopically labeled protein and antibiotic laced M9 minimal media. Cells were grown at 37°C until an OD_600_ of 0.6-0.9, induced with 0.4 mM isopropyl-1-thio-d-galactopyranoside (IPTG), and shaken overnight at 21°C and 180 rpm. Cells were harvested by centrifugation and lysed by sonication in low-imidazole binding buffer (**Table S3**) with 1mM PMSF and clarified by centrifugation. Clarified lysate was loaded onto an equilibrated HisTrap FF column (Cytiva), washed with five column volumes, and released with high-imidazole buffer (**Table S3**). ARC4 underwent an additional overnight cleavage with PreScission protease, and an additional His-trap purification to remove the GST-tag. All three proteins underwent a final preparatory step and buffer exchange with size-exclusion chromatography (Column 2) into NMR Buffer (**Table S3**).

### NMR Experiments and Data Analysis

All experiments were conducted at 5°C on a Bruker 600 MHz Avance Neo with a cryoprobe. The ^15^N-^1^H heteronuclear single-quantum correlation (HSQC) spectra for the titration were acquired with a nonuniform sampling (NUS) scheme generated by the NUS@HMS scheme generator software ^60^ with 1,024 complex data points in the direct dimension and 50% sampling of the original 200 real points in the indirect ^15^N dimension, and 32 scans. The recycle delay was 1.3 s, and the spectral widths were 13 and 35 ppm for the ^1^H and ^15^N dimensions, respectively. The time-domain data were processed using NMRPipe ^61^ and analyzed using CcpNmr ^62^ and POKY ^63^. HSQC of the partially assigned N.V. SPIDER was overlaid with the assigned HSQC of S519N to confirm the chemical shift ^31^. The chemical shift perturbations (CSPs) were monitored by overlaying ^15^N-^1^H HSQC spectra of apo ^15^N-labeled SPIDER and in 1:1, 1:2, 1:4, 1:10, and 1:12 mixtures with unlabeled ARC4 using the following equation:

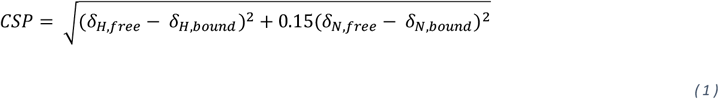

Where *δ*_*H*_ and *δ*_*N*_ are the chemical shifts of a peak in the ^1^H and ^15^N dimensions, respectively.

In addition to CSPs, peak broadening was quantified by calculating the ratio of the peak intensities of apo SPIDER and SPIDER bound to ARC4 at different concentration ratios.

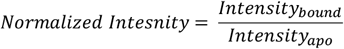

### Microscale thermophoresis (MST)

Microscale thermophoresis (MST) was performed on a NanoTemper Monolith NT.115 Pico instrument (NanoTemper Technologies GmbH) at 25°C using auto-detect Pico Red at 20% excitation power. ARC4 was fluorescently labelled by incubating 100 *μ*l of 2 *μ*M protein solution with 100 *μ*l of 6 *μ*M red fluorescent dye NT-647 Maleimide for 30 min. The reaction mixture was centrifuged for 10 min at 4°C and 15,000 g speed and dialyzed overnight against the NMR buffer mentioned above. 10 nM of the labeled ARC4 and 16 two-fold dilution series of SPIDER were loaded into sixteen premium capillaries (NanoTemper Technologies GmbH; highest concentrations were 195 *μ*M of SPIDER N.V., and 200 *μ*M of SPIDER S519N). The sigmoidal curves obtained were analyzed to extract the *K*_D_.

## Statistical Analyses

One-Way Analysis of Variance (ANOVA) and Tukey post-hoc test for the sphere forming and Aldefluor assays, and Fisher’s exact test for the onset of hair graying were done with GraphPad Prism 8 software. All comparisons were two-sided, with *p-values* < 0.05 considered significant.

### RNA-seq analyses with SASH1 knockdown in human primary melanocytes in sphere cultures

1.5 × 10^6^ hEMnMP cells were transfected with 100 pmol siRNAs with a Nucleofector 4D system, using the P2 solution and program DS-137. Three nucleofections were transferred to 6 cm dishes and allowed to recover in medium 254 + HMGS-2 for 5 hours. Cells were detached, and 1 × 10^6^ cells were suspended in 25 ml of stem cell medium and seeded in 2 × 10 cm PolyHEMA coated dishes, as in ^24^. After 19 hrs in sphere conditions, floating cells were collected and rinsed once with PBS, followed by total RNA extraction with the Qiagen RNeasy Plus Mini kit, including the Qiashredder column.

Triplicate RNAs were sent to the University of Colorado Anschutz Medical Campus Genomics and Microarray Core for RNA library construction and sequencing, as in ^64^. FASTQ files generated were processed using the Cufflinks/Cuffdiff RNAseq workflow and aligned to Ch37/hg19 ^65^. Cuffdiff analysis generated fragment per kilobase per million mapped read (FPKM) values and performed differential gene expression analysis between sample groups. The fragment per kilobase per million (FPKM) values were analyzed by Gene Set Enrichment Analysis (GSEA, Broad Institute, v6.2MSigDB) using Hallmark pathway as gene sets ^66,67^.

## Notes

### Competing Interest Statement

The authors have declared no competing interest.

