## Supplemental materials for "SASH1 interacts with TNKS2 and promotes human melanocyte stem cell maintenance"

### Supplementary Materials

#### Table of Contents

**Figure S1.** Stem-like characteristics in human primary melanocytes lost with SASH1 knockdown

**Figure S2**. RT-PCR validation of siTNKS2 knockdown for the sphere assays in Fig.3D

**Figure S3.** 1D $^{1}$H-NMR spectra used to confirm the proper folding of recombinant proteins

**Figure S4**. Select peak overlays showing differences between N.V. and S519N behavior upon titration of ARC4

**Figure S5.** SASH1 is involved in mitosis

**Table S1.** Antibodies and Magnetic Beads

**Table S2.** Vectors

**Table S3.** Buffers and Media

**Table S4.** Chemicals and Reagents

**Table S5.** Preparatory Columns

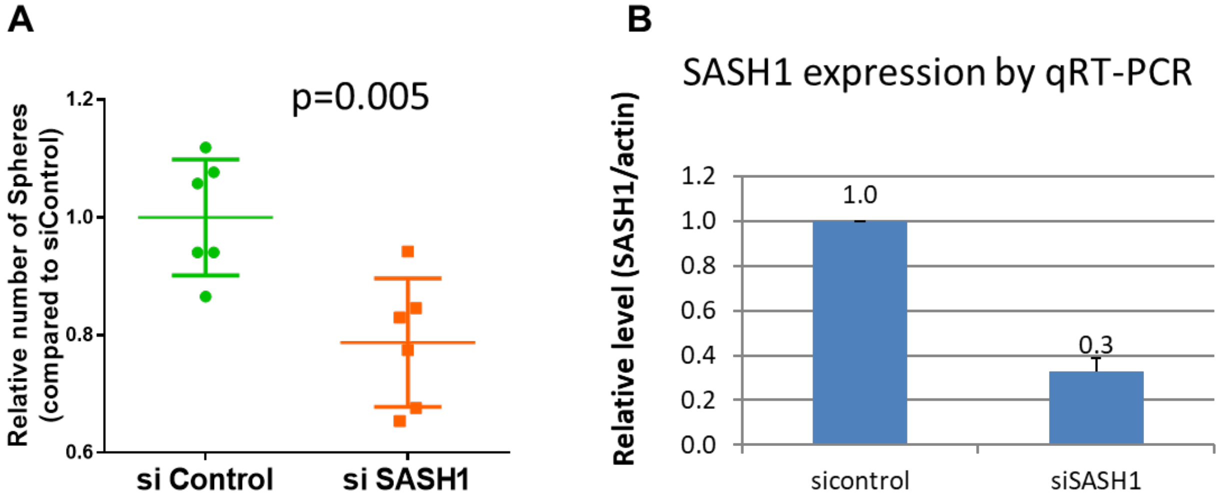


**Figure S1. Knocking down of SASH1 decreased stem-like characteristics in human primary melanocytes.** Human primary melanocyte cells were transfected with control siRNA (sicontrol) or siRNA against SASH1 (siSASH1). Cells were then subjected to the sphere forming assay (A), or quantitative RT-PCR of SASH1 normalized by Actin (B).

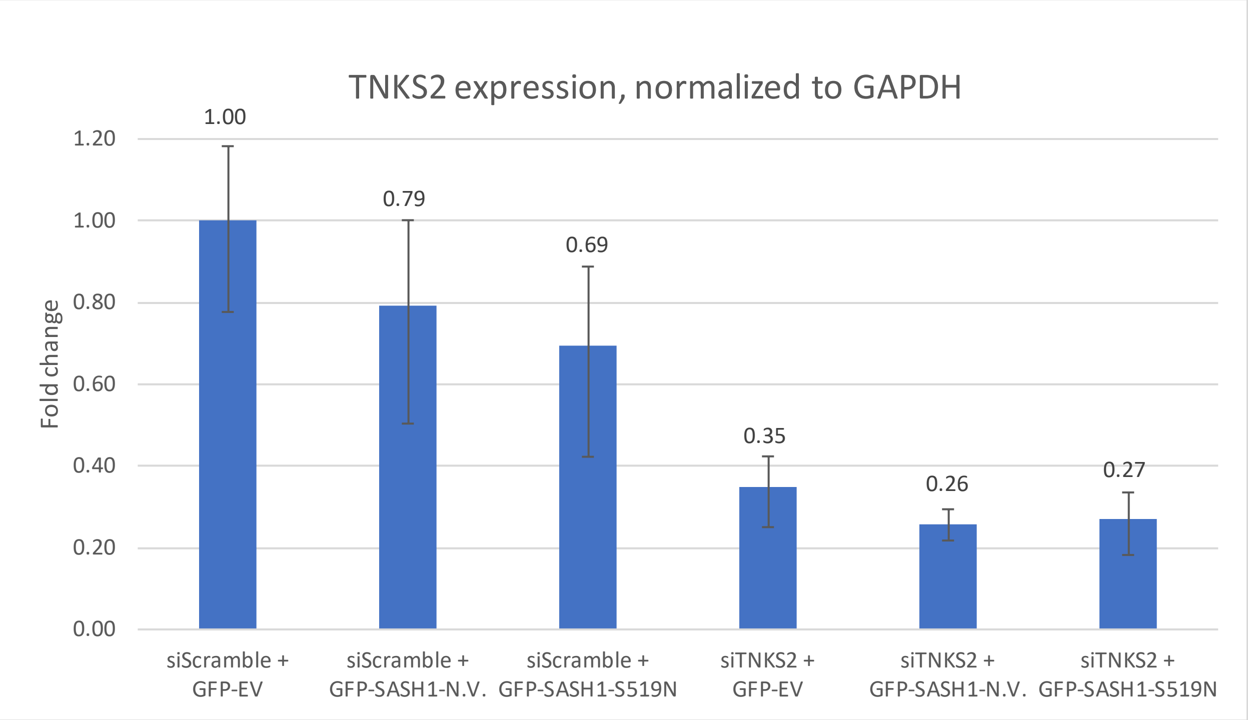


**Fig. S2**. **RT-PCR validation of siTNKS2 knockdown for the sphere assays in Fig.3D**. Human primary MCs were transfected with expression vectors of SASH1 (control vector, EV; Non-variant, N.V. or S519N SASH1), with siRNAs against TNKS2 (siTNKS2) or scramble control (siScramble). Cells were then subjected to sphere forming assays. Total RNA was harvested at 5 hr post transfection, prior to seeding in the sphere assays.

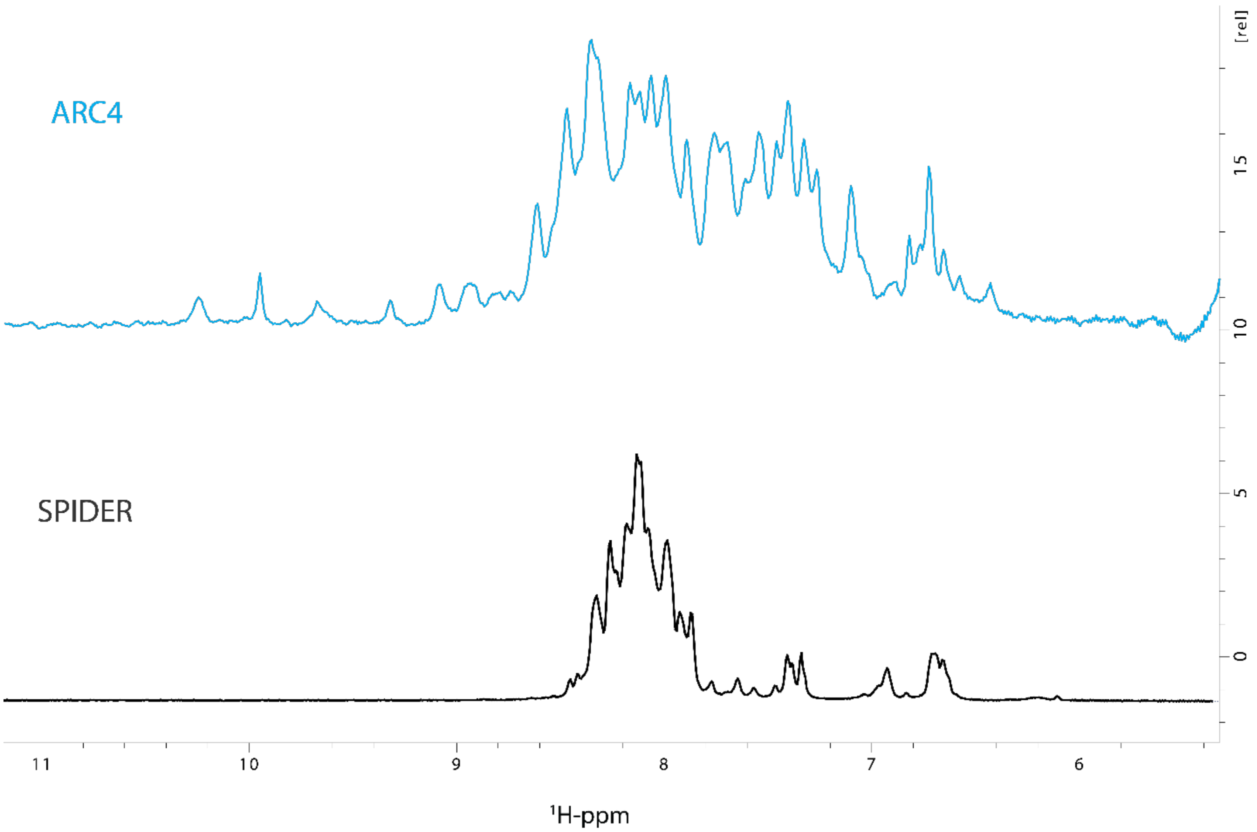


**Figure S3. 1D $^1$H-NMR spectra used to confirm the proper folding of recombinant protein ARC4.** 1D $^1$H-NMR spectrum TNKS2-ARC4 (cyan) shows clear peak dispersion in the amide-backbone region of over 2 ppm (~7.1 – 9.2), confirming a well folded protein. The peaks of the backbone-amide region of N.V. SPIDER (black) are < 1 ppm (~7.8 – 9.4) indicative of a disordered protein. Both spectra were obtained on a Bruker 600 MHz spectrometer.

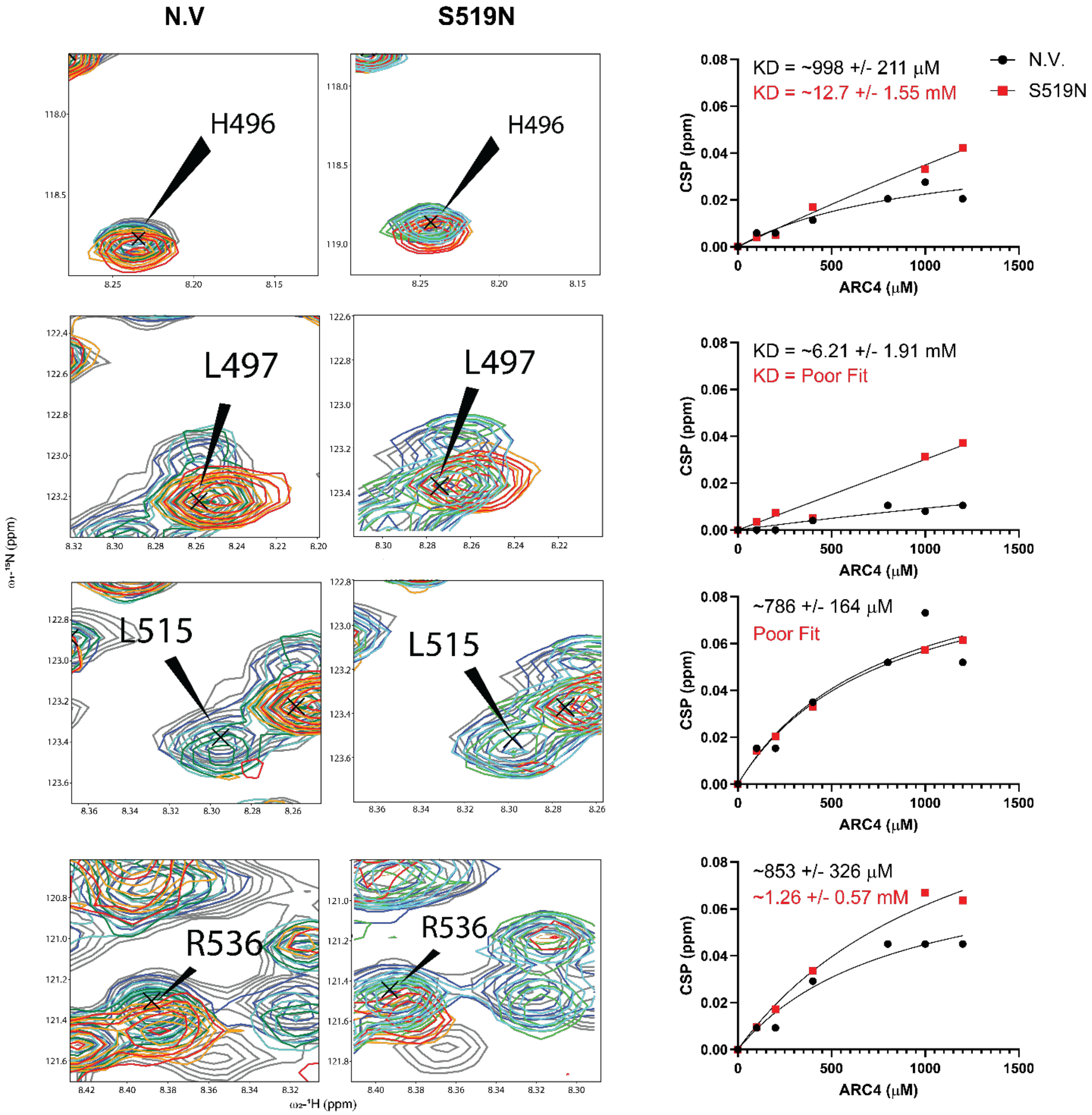


**Figure S4. Select peak overlay showing differences between N.V. and S519N in their behaviors upon titration of ARC4.** $K_D$ traces with fitted values of both N.V. and S519N (black and red, respectively) have been overlaid adjacent to the CSPs. Insets display atomic level changes in binding kinetics.

**A**

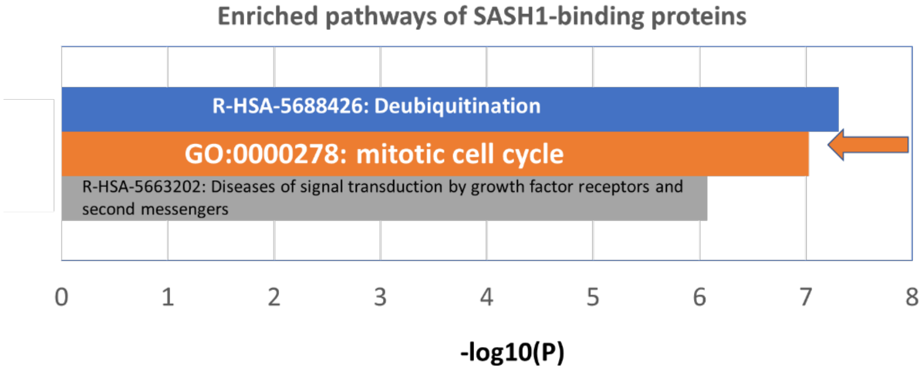


**B**

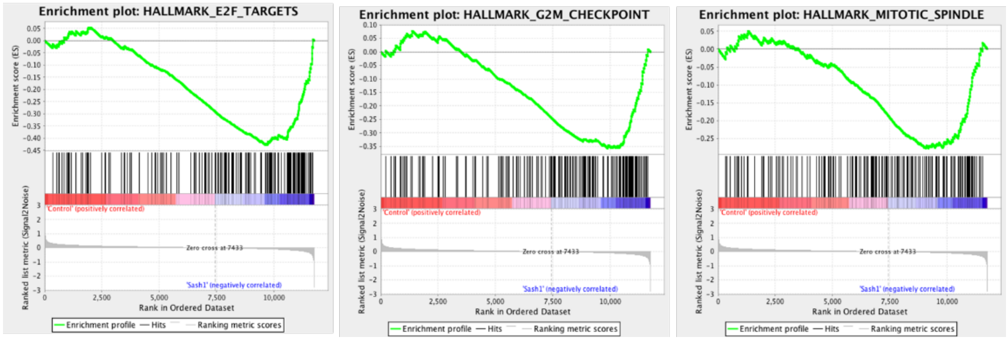


**Figure S5. SASH1 is involved in mitosis.** (A) The top enriched pathways/bioprocesses for SASH1-binding proteins with –log10 (p value) > 6, indicated in X-axis. Gene lists of three Y2H screens, with full length SASH1 as bait, were run on Metascape.org for the enrichment analyses. The mitotic cell cycle is a top pathway. (B) Knockdown of SASH1 induced G2/M and mitotic spindle checkpoints in human primary melanocytes. Human primary MCs were transfected with siControl (Control) or siSASH1 (SASH1), cultured in sphere conditions, and subjected to RNAseq analyses. siSASH1 decreased SASH1 expression by ~60%. GSEA enrichment plots for the top differentiated expressed Hallmark pathways are shown here: E2F targets (p-value = 0.0001, FDR = 0.002), G2M Checkpoint (p = 0.004, FDR = 0.031), Mitotic spindle (p = 0.058, FDR = 0.12). Top panel: running enrichment scores in green. Bottom panel: black vertical bars show position of gene set members within the ranked list.

#### Table S1: Antibodies and Magnetic Beads

| | Target | Application | Host | Company | Part Number | Dilution | Developer |
|---|---|---|---|---|---|---|---|
| 1 | HA-tag beads | CoIP, magnetic beads | Mouse IgG1 | Thermo Scientific | 88836 | 1:16 beads:lysate | N/A |
| 2 | HA-tag | WB Primary Antibody | Rabbit | Cell Signaling Technology | 3724 | 1:50000 | Super Signal West Pico Plus Chemiluminescent Substrate (Thermo 34580) |
| 3 | TNKS1/2 | WB Primary Antibody | Mouse | Santa Cruz | 365897 | 1:2000 | Super Signal West Pico Plus Chemiluminescent Substrate (Thermo 34580) |
| 4 | GAPDH | WB Primary Antibody | Rabbit | Cell Signaling Technology | 5174 | 1:1000 | Super Signal West Femto Maximum Sensitivity Substrate (Thermo 34095) |
| 5 | Tubulin beta | WB Primary Antibody | Rabbit | Thermo Scientific | RB-9249-P0 | 1:5000 | |
| 6 | Rabbit-HRP | WB Secondary Antibody | Goat | Cell Signaling Technology | 7074 | 1:10000 | N/A |
| 7 | Mouse-HRP | WB Secondary Antibody | Goat | Cell Signaling Technology | 7076 | 1:10000 | N/A |
| 8 | Ki-67 | Immunofluorescence Primary Antibody | Mouse IgG1 | BD Pharmingen | 556003 | 1:40 | N/A |
| 9 | Tyrosinase [T311] | Immunofluorescence Primary Antibody | Mouse IgG2a | Abcam | ab738 | 1:10 | N/A |
| 10 | MouseIgG1-AlexaFluor-594 | Immunofluorescence Secondary Antibody | Goat | Invitrogen | A21125 | 1:250 | N/A |
| 11 | MouseIgG2a-AlexaFluor-488 | Immunofluorescence Secondary Antibody | Goat | Invitrogen | A21131 | 1:250 | N/A |

#### Table S2: Vectors

| | Protein of Interest | Protein Region | Vector | Antibiotic | N-terminal Tags | C-terminal Tags | Supplier |
|---|---|---|---|---|---|---|---|
| 1 | TNKS2-ARC4 | 488-649 | pGEX-6P-1 | Carbenicillin | GST Tag, PreScission Cleavage Site | 6x-Histadine Tag | GenScript |
| 2 | SASH1-SPIDER (N.V and S519N) | 396-546 | pET-28a (+) | Kanamycin | 6x-Histadine Tag, thrombin cleavage site | N/A | GenScript |
| 3 | pMEV-2HA (Empty Vector) | N/A | N/A | Kanamycin | 2x HA tag | N/A | BioMyx #P1001 |
| 4 | SASH1 (N.V.) | full length | pMEV-2HA | Kanamycin | 2x HA tag | N/A | OriGene #TC100438 |
| 5 | SASH1-(S519N) | full length | pMEV-2HA | Kanamycin | 2x HA tag | N/A | Site directed mutagenesis with Agilent Quick Change II |
| 6 | pEZYflag (Empty Vector) | N/A | N/A | Chloramphenicol, Ampicillin | FLAG tag | N/A | Addgene #18700 |
| 7 | TNKS2 | full length | pFLAG | Ampicillin | FLAG tag | N/A | Addgene #34691 |

#### Table S3: Buffers and Media

| | Buffer | Buffer Contents | pH |
|---|---|---|---|
| 1 | Low Imidazole-Binding | 20 mM HEPES, 200 mM NaCl, 1 mM EGTA, 1 mM $MgCl_2$, 1 mM $NaN_3$, 20 mM imidazole | 7.3 |
| 2 | High Imidazole-Elution | 20 mM HEPES, 200 mM NaCl, 1 mM EGTA, 1 mM $MgCl_2$, 1 mM $NaN_3$, 200 mM imidazole | 7.3 |
| 3 | Size exclusion and NMR Buffer | 20 mM NaP, 100 mM NaCl, 1 mM DTT, 0.02% $NaN_3$ | 7.0 |
| 4 | Co-Immunoprecipitation Lysis/Wash Buffer | 137 mM NaCl, 20 mM Tris-HCl, 1% Triton X-100, 1% Glycerol, 1 mM EDTA, 2.5 uM PDD 00017273, 2X HALT Protease/Phosphatase inhibitors without EDTA, 1 mM PMSF | 8.0 |
| 5 | Harsh western blot stripping buffer (Abcam's) | 62.5 mM Tris-HCl, 2% SDS, 0.8% betamercaptoethanol | 6.8 |
| 6 | 293FT culture medium | High Glucose DMEM, 1 mM Sodium Pyruvate, 4 mM L-Glutamine, 10% FBS | |
| 7 | CoIP elution buffer | 4X Laemmli buffer, 10% betamercaptoethanol | 6.8 |
| 8 | Western blot transfer Buffer | 1X Tris-Glycine Buffer, 20% Methanol, 0.05% SDS | 8.3 |
| 9 | Western blot blocking buffer | 5% non-fat milk in TBS | 7.5 |
| 10 | Stem/Sphere culture medium | DMEM/F12, 2% B-27, 2X Pen/Strep, 40 pg/ml bFGF, 4 pg/ml EGF, 480 pg/ml Heparin | |

##### Table S4: Chemicals and Reagents

| | Chemical/Reagent Name | Company | Product Number | Additional information |
|---|---|---|---|---|
| 1 | 293FT mammalian cells | Life Technologies | R70007 | |
| 2 | High Glucose DMEM | Cytiva Hyclone | SH3024301 | |
| 3 | Glutamax I | Gibco | 35050061 | |
| 4 | Fetal Bovine Serum | Peak | PS-FB3 | |
| 5 | Effectene | Qiagen | 301425 | |
| 6 | 4-12% Tris-Glycine SDS PAGE gel | Invitrogen | WXP41226BOXA | |
| 7 | 10X Tris-Glycine SDS Buffer | Bio-rad | 1610732 | |
| 8 | 10X Tris-Glycine Buffer | Bio-rad | 1610734 | |
| 9 | Ponceau S Staining Solution | Cell Signaling Technology | 59803 | |
| 10 | siRNA against SASH1 | Sigma | Custom Mission siRNA | 5'-->3': CAACAUUGCCUUUAAUGAA UUCAUUAAAGGCAAUGUUG |
| 11 | hEMnMP: human Epidermal Melanocytes, neonatal, Moderately Pigmented | Gibco | C1025C | |
| 12 | Medium 254 | Gibco | M254500 | |
| 13 | HMGS-2: Human Melanocyte Growth Supplement 2 | Gibco | S0165 | |
| 14 | Allstars negative control siRNA | Qiagen | SI03650318 | |
| 15 | Amaxa P2 Primary Cell 4D-Nucleofector X Kit | Lonza | V4XP-2012 | |
| 16 | OptiMEM | Gibco | 31985062 | |
| 17 | Aldefluor kit | Stem Cell Technologies | #01700 | |
| 18 | qPCR primer pair, SASH1 | Integrated DNA Technologies | custom | 5'→3'<br>F: ATGGCTCTCCTGAGAAACCTC<br>R: CTGAAGGTTTTGACCAACTTCC |
| 19 | qPCR primer pair, ACTIN | Integrated DNA Technologies | custom | 5'→3'<br>F: GACAGGATGCAGAAGGAGATTACT<br>R: TGATCCACATCTGCTGGAAGGT |

##### Table S5: Preparatory Columns

| | Column Name | Packing | Dimensions | Volume | Supplier | Part Number |
|---|---|---|---|---|---|---|
| 1 | HisTrap FF | Ni Sepharose 6 Fast Flow | 5 x 1 mL | 5mL | Cytiva | 17531901 |
| 2 | HiLoad 16/600 Superdex 75 pg | Superdex 75 prep grade resin | 16 x 600 mm | 120mL | Cytiva | 28989333 |
